# Evaluation of six phase encoding based susceptibility distortion correction methods for diffusion MRI

**DOI:** 10.1101/766139

**Authors:** Xuan Gu, Anders Eklund

## Abstract

**Purpose:** Susceptibility distortions impact diffusion MRI data analysis and is typically corrected during preprocessing. Correction strategies involve three classes of methods: registration to a structural image, the use of a fieldmap, or the use of images acquired with opposing phase encoding directions. It has been demonstrated that phase encoding based methods outperform the other two classes, but unfortunately, the choice of which phase encoding based method to use is still an open question due to the absence of any systematic comparisons.

**Methods:** In this paper we quantitatively evaluated six popular phase encoding based methods for correcting susceptibility distortions in diffusion MRI data. We employed a framework that allows for the simulation of realistic diffusion MRI data with susceptibility distortions. We evaluated the ability for methods to correct distortions by comparing the corrected data with the ground truth. Four diffusion tensor metrics (FA, MD, eigenvalues and eigenvectors) were calculated from the corrected data and compared with the ground truth. We also validated two popular indirect metrics using both simulated data and real data. The two indirect metrics are the difference between the corrected LR and AP data, and the FA standard deviation over the corrected LR, RL, AP and PA data.

**Results:** We found that *DR-BUDDI* and *TOPUP* offered the most accurate and robust correction compared to the other four methods using both direct and indirect evaluation metrics. *EPIC* and *HySCO* performed well in correcting *b*_0_ images but produced poor corrections for diffusion weighted volumes, and also they produced large errors for the four diffusion tensor metrics. We also demonstrate that the indirect metric (the difference between corrected LR and AP data) gives a different ordering of correction quality than the direct metric.

**Conclusion:** We suggest researchers to use *DR-BUDDI* or *TOPUP* for susceptibility distortion correction. The two indirect metrics (the difference between corrected LR and AP data, and the FA standard deviation) should be interpreted together as a measure of distortion correction quality. We also suggest that indirect metrics must be interpreted cautiously when evaluating methods for correcting susceptibility distortions in diffusion MRI data.

## 1 INTRODUCTION

Analysis of diffusion MRI data is confounded by the presence of susceptibility distortions, caused by an off-resonance field induced by differences in magnetic susceptibility at the air-tissue interface. There are a number of techniques available for correcting susceptibility distortions. Broadly, these techniques can be divided into three types: registration based (RB) methods, fieldmap based (FB) methods and phase encoding based (PB) methods. The first approach involves registration of the distorted image to a structural image without distortions. The second approach involves estimating a map of the *B*_0_ inhomogeneity, and using this along with information about the diffusion acquisition protocol to correct for the distortions. The third approach is based on estimating the underlying distortions using additional data acquired with a different phase encoding direction. For example, it is common to collect LR (left right) and RL (right left) images, or AP (anterior posterior) and PA (posterior anterior) images. Phase encoding based techniques have been demonstrated to outperform the other two approaches (Graham et al., 2017; Esteban et al., 2014), at the cost of a longer scan time.

There are many software packages providing phase encoding based tools for correcting susceptibility distortions, e.g., *animaDistortionCorrection* (aDC) (Voss et al., 2006), *animaBMDistortionCorrection* (aBMDC) (Hedouin et al., 2017), *DR-BUDDI* (Irfanoglu et al., 2015), *EPIC* (Holland et al., 2010), *HySCO* (Ruthotto et al., 2013) and *TOPUP* (Andersson et al., 2003), summarized in Table 1. To date, there is no systematic comparison of existing phase encoding based methods for susceptibility distortion correction. See Table 2 for an overview of previous comparisons of different distortion correction tools.

**TABLE 1.**
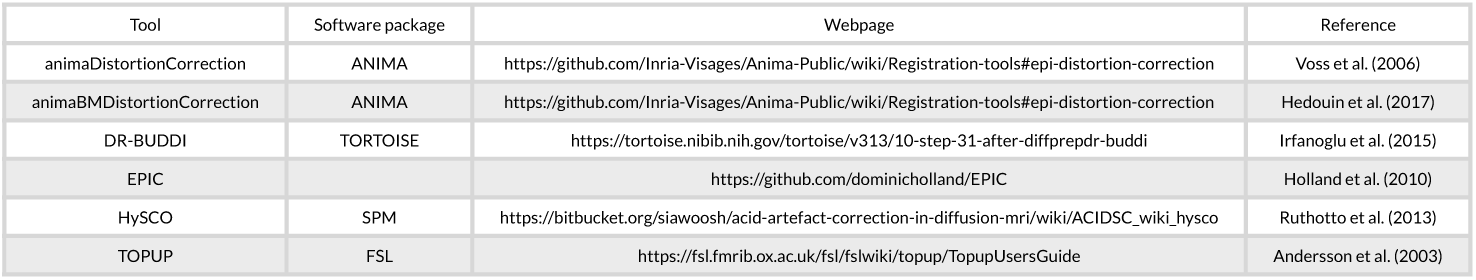
Phase encoding based susceptibility distortion correction tools evaluated in this paper.

**TABLE 2.**
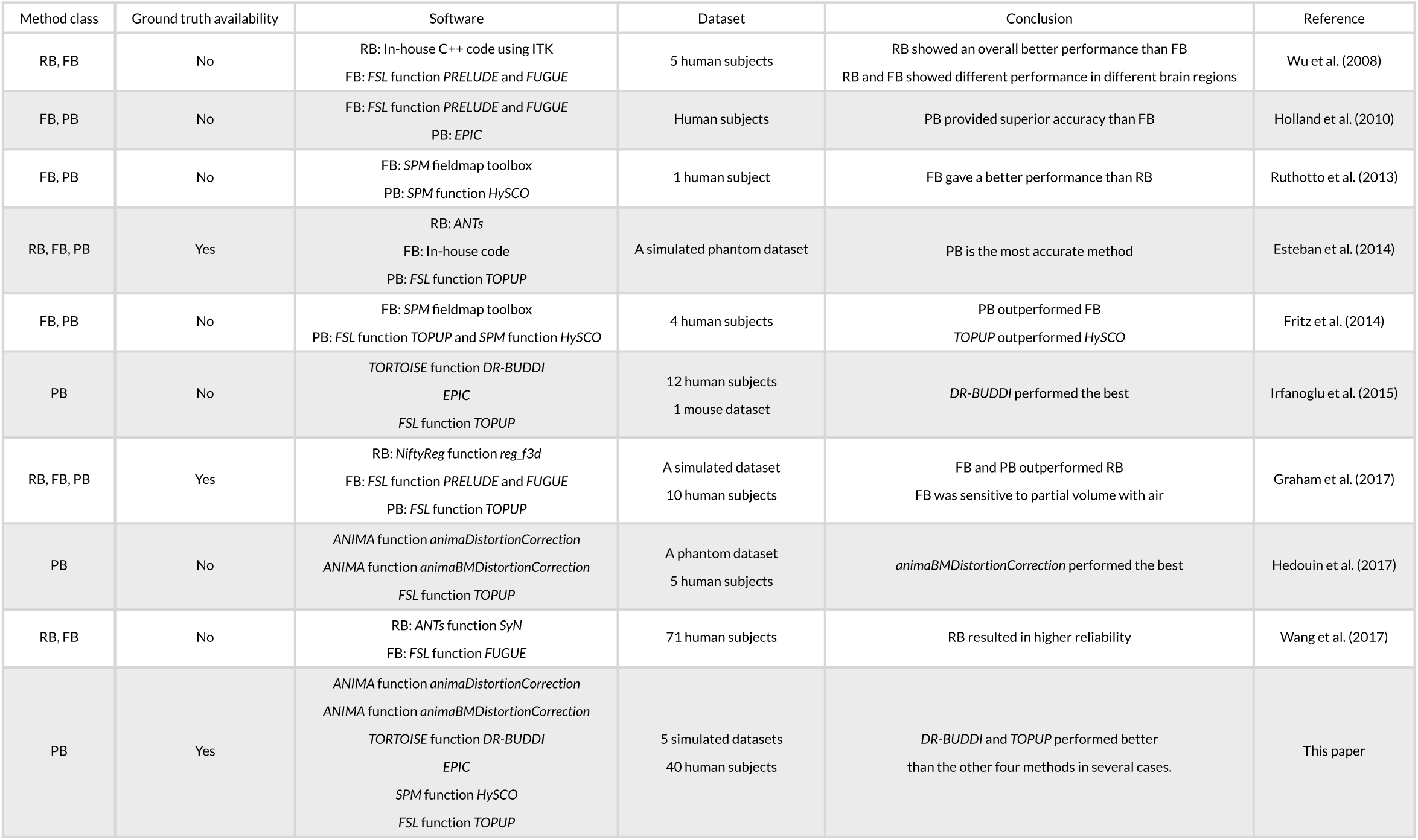
A list of susceptibility distortion correction evaluation papers. RB: registration based methods, FB: fieldmap based methods, PB: phase encoding based methods.

The lack of ground truth means that evaluations are typically indirect or qualitative (Jezzard and Balaban, 1995; Wu et al., 2008; Bhushan et al., 2012; Ruthotto et al., 2013; Fritz et al., 2014; Irfanoglu et al., 2015; Taylor et al., 2016; Hedouin et al., 2017; Wang et al., 2017; Irfanoglu et al., 2018). Only a few investigations have been carried out with the presence of a ground truth for evaluation of susceptibility distortion correction (Andersson et al., 2003; Esteban et al., 2014; Graham et al., 2017). Hedouin et al. (2017) compared *aDC, aBMDC* and *TOPUP* using phantom data and human data. For the phantom data, both the *aBMDC* and *TOPUP* corrected images appear visually similar for correcting the distortion, while *aDC* gives visually poorer results. *aBMDC* outperformed *aDC* and *TOPUP* by obtaining smaller landmark position errors. For human data, images corrected using *aDC* contain a mismatch around the lateral ventricles compared with respect to a structural (T1-weighted) image. *aBMDC* and *TOPUP* both obtain a corrected image very close to the structural T1-weighted image. *aBMDC* and *TOPUP* show a very high similarity between the two corrected images C_AP+PA_ and C_LR+RL_, outperforming *aDC*. Irfanoglu et al. (2015) compared *EPIC, DR-BUDDI* and *TOPUP* using human data. *DR-BUDDI* produced sharper images than *EPIC* and *TOPUP*, showing clearly visible tissue interfaces. Areas such as the inferior temporal lobes and the olfactory bulbs were more accurately reconstructed by *DR-BUDDI* than *EPIC* and *TOPUP. DR-BUDDI* resulted in the lowest variability between the two corrected images C_AP+PA_ and C_LR+RL_, followed by *TOPUP* and then *EPIC*. Overall, *DR-BUDDI* corrected images showed a higher correlation with the undistorted T2-weighted image than did *EPIC* and *TOPUP*.

In this work, we undertake a comparison of six phase encoding based methods for susceptibility distortion correction using both simulated diffusion data and real diffusion data, see Table 2 for differences between our study and previous comparisons. A brief summary of the six methods is as follows.

- *aDC*: The *aDC* method estimates the displacement field based on the image cumulative distribution function along each line in the PE direction. This method is rather sensitive to noise and it realigns every line independently, so in order to reduce the effect of noise and increase the coherence between the corrected lines, a 3D Gaussian smoothing is applied on the estimated displacement field, which leads to a trade-off between regularity and precision.
- *aBMDC*: The *aBMDC* method adopts the symmetric diffeomorphic image registration idea from Avants et al. (2008) to make the transformed LR (or AP) and RL (or PA) images as similar as possible. The transformation field is obtained using a block-matching algorithm (Ourselin et al., 2000). The squared correlation coefficient is used as the similarity measure between blocks.
- *DR-BUDDI*: The *DR-BUDDI* method also adopts the symmetric diffeomorphic image registration idea from Avants et al. (2008). The first part of the cost function is designed to maximize the similarity between the structural image and the transformed LR/RL (or AP/PA) images. The second part of the cost function is designed to maximize the similarity between the structural image and the geometric average of the transformed LR and RL images. The transformation is constrained along the PE direction. Two co-dependent transformations (one for mapping LR to RL (or AP to PA), the other for mapping RL to LR (or PA to AP)) are used to account for differences in the inhomogeneity between LR (or AP) and RL (or PA) acquisitions. Unlike other methods, *DR-BUDDI* estimates the transformation not only for *b*_0_ images but based on a weighted sum over different diffusion weighted images, in order to preserve tract structures.
- *EPIC*: The *EPIC method* estimates the displacement field by minimizing the sum of squared differences between the transformed LR (or AP) and RL (or PA) images. Two regularization terms are used to regularize the smoothness and amplitude of the displacement. The undistorted image can then be restored using the estimated displacement field.
- *HySCO*: The *HySCO* method adopts the physical distortion model from Chang and Fitzpatrick (1992) and minimizes a distance function to estimate the inhomogeneity, such that the transformed LR (or AP) and RL (or PA) images become as similar as possible. Two regularization terms are used to ensure a differentiable inhomogeneity and positive intensity modulations.
- *TOPUP*: The *TOPUP* method models the displacement field as a linear combination of basis warps consisting of a truncated 3D cosine transform. The weights of the basis warps are estimated using an iterative procedure by minimizing the sum of squared differences between the transformed LR (or AP) and RL (or PA) images. Two options are available for obtaining the distortion free image, they are based on least-squares and Jacobian modulation. The resolution of the displacement field is limited by the highest frequency component of the cosine transform.

We used the POSSUM (Drobnjak et al., 2006, 2010) based diffusion MRI simulator (Graham et al., 2016, 2017), in order to produce realistic diffusion data with susceptibility distortions typically seen in real data. Simulated data can provide ground truth that enables direct and quantitative evaluation. Our analysis directly measures the ability to correctly recover distortion-free data by comparing the corrected *b*_0_ image with its ground truth. We also investigate the suitability of two commonly used indirect metrics, i.e. the difference of the corrected data from the LRRL and APPA pairs (Ruthotto et al., 2013; Graham et al., 2017), and the FA standard deviation over the corrected LR, RL, AP and PA data (Wu et al., 2008; Irfanoglu et al., 2015). We hope that this work will enable researchers to make more carefully informed choices when designing their processing pipelines.

## 2 DATA

### 2.1 Simulated data

The diffusion data was simulated with 11 volumes of *b* = 700 s/mm^2^, 12 volumes of *b* = 2000 s/mm^2^ and 1 volume of *b* = 0. The input to the POSSUM diffusion MRI simulator is a collection of three 3D anatomical volumes: grey matter, white matter and cerebro-spinal fluid (CSF). The voxel values in these segmentations reflect the proportion of tissue present in each voxel, in the range [0,1]. The input was generated from the T1-weighted structural image from HCP using the *FSL* function *FAST* (Zhang et al., 2001). The representation of diffusion weighting was achieved by a spherical harmonic fit of order *n* = 8 to the *b* = 1000 s/mm^2^ shell of the diffusion data, using the *Dipy* (Garyfallidis et al., 2014) *module reconst.shm*. The MR parameters used are listed in Table 3. We used a matrix size of 72 × 86, 55 slices and a voxel size of 2.5 mm isotropic. The TE was 109 ms, the TR was 9.15 s and the flip-angle was 90 °. The PE bandwidth per pixel was 17.1 Hz for LR and RL data, and 14.3 Hz for AP and PA data. The fieldmap was generated from one phase difference volume and two magnitude volumes (one for each echo time) from HCP using the *FSL* function *fsl_prepare_fieldmap* (Jenkinson, 2003). To generate a tight brain extration for *fsl_prepare_fieldmap*, the brain mask created by the *FSL* function *BET* (Smith, 2002) was further eroded using a 5 mm box kernel. The generated fieldmap was linearly registered to the T1-weighted structural image using the *FSL* function *FLIRT* (Jenkinson and Smith, 2001; Jenkinson et al., 2002). Diffusion data was simulated with four PE directions, i.e., left-right (LR), right-left (RL), anterior-posterior (AP) and posterior-anterior (PA). No other distortions (e.g. eddy-currents and head motion) were included in the simulations. We also simulated a ground truth set, acquired with the same acquisition parameters but no input susceptibility fieldmap. We simulated diffusion data for five subjects (100206, 100307, 100408, 100610, 101006) from the Human Connectome Project (HCP)^1^ (Van Essen et al., 2013; Glasser et al., 2013).

**TABLE 3.** A list of MR parameters (relaxation times T1, T2*, spin density, and chemical shift value) used in our simulations within POSSUM.

|  | T1 (ms) | T2* (ms) | Spin density | Chemical shift | T2 (ms) |
| --- | --- | --- | --- | --- | --- |
| GM | 1331 | 51 | 0.86 | 0 | 75 |
| WM | 832 | 44 | 0.77 | 0 | 70 |
| CSF | 3700 | 500 | 1 | 0 | 500 |

### 2.2 Real data

We used 40 subjects from the developing HCP project (Hughes et al., 2017; Bastiani et al., 2019). It provides diffusion data acquired with four PE directions: AP, PA, LR and RL, enabling evaluation using the indirect metric (i.e. comparing APPA corrected to LRRL corrected). The data was acquired on a 3T Philips Achieva scanner and consists of 4 shells: 20 volumes of *b* = 0, 64 volumes of *b* = 400 s/mm^2^, 88 volumes of *b* = 1000 s/mm^2^ and 128 volumes of *b* = 2600 s/mm^2^. The data was acquired using TR=3800 ms and TE=90 ms. The matrix size is 128 × 128, the number of slices is 64 and the acquired voxel size is 1.17 × 1.17 × 1.5 mm.

## 3 METHODS

For the simulated data, we used the *FSL* function *BET* (Smith, 2002) to create the brain mask from the distortion-free *b*_0_ image. Diffusion tensor fitting and FA calculation were performed using the *FSL* function *dtifit*. For simulated data, we evaluated the ability of each method to recover the correct intensity at each voxel, by computing error maps between the distortion corrected data and ground truth. We also investigated two indirect metrics. One is the difference of the corrected data from the LRRL and APPA pairs (Ruthotto et al., 2013; Graham et al., 2017), and the other is the FA standard deviation over the corrected LR, RL, AP and PA data (Wu et al., 2008; Irfanoglu et al., 2015). Comparing the uncertainty of data from different preprocessing pipelines is a way to determine if one pipeline is better than the other (Sjölund et al., 2018; Gu et al., 2019). For the real data, we used the brain mask provided with the dataset. We corrected for head motion between LR, RL, AP and PA scans by registering all volumes to the first volume using the *FSL* function *FLIRT* (Jenkinson and Smith, 2001; Jenkinson et al., 2002). For real data, we used indirect evaluation. We compared the corrected data from the LRRL pair with the result from the APPA pair, since ideally the corrected results from the two pairs should be the same.

For each susceptibility distortion correction tool we used default settings and steps provided by the software’s basic help documentation unless otherwise stated. For *DR-BUDDI*, we used the command *DR_BUDDI_withoutGUI* because it is faster than *DR_BUDDI*. By default, the first step in *DIFFPREP* performs Gibbs ringing correction, denoising, head motion correction and eddy-current distortion correction, prior to the second step susceptibility distortion correction by *DR-BUDDI*. However, the simulated data in this paper have only susceptibility distortions, corrections by *DIFFPREP* are therefore not necessary and can even be counterproductive. Additionally, the use of a structural image in *DR-BUDDI* will introduce deviation from the ground truth due to the registration between diffusion and structural images. Therefore, we ran *DR-BUDDI* for the simulated data without the corrections from *DIFFPREP* and without a structural image. For the real data, we ran *DR-BUDDI* with the default setting together with *DIFFPREP*. There is no method referred to in any of the *EPIC* documentation for choosing an alternate phase direction than the *y*-direction (or the LRRL direction with regards to this study). We manually rotated the LR and RL data 90 degrees in the *x* − *y* plane before feeding them to *EPIC*. The data was finally rotated 90 degrees in the other direction after correction. We share our processing scripts on Github ^2^, such that other researchers can reproduce and extend our findings (Eklund et al., 2017).

## 4 RESULTS

### 4.1 Simulated data

Figure 1 shows the simulated diffusion data and the fieldmap for HCP Subject 100206, the simulated distortions look very realistic. We investigated how different levels of distortion affect the correction performance by simulating three levels of susceptibility distortion, as shown in Figure 2. The distortion level was controlled by dividing the fieldmap by a factor 1, 2 or 4.

**FIGURE 1.**
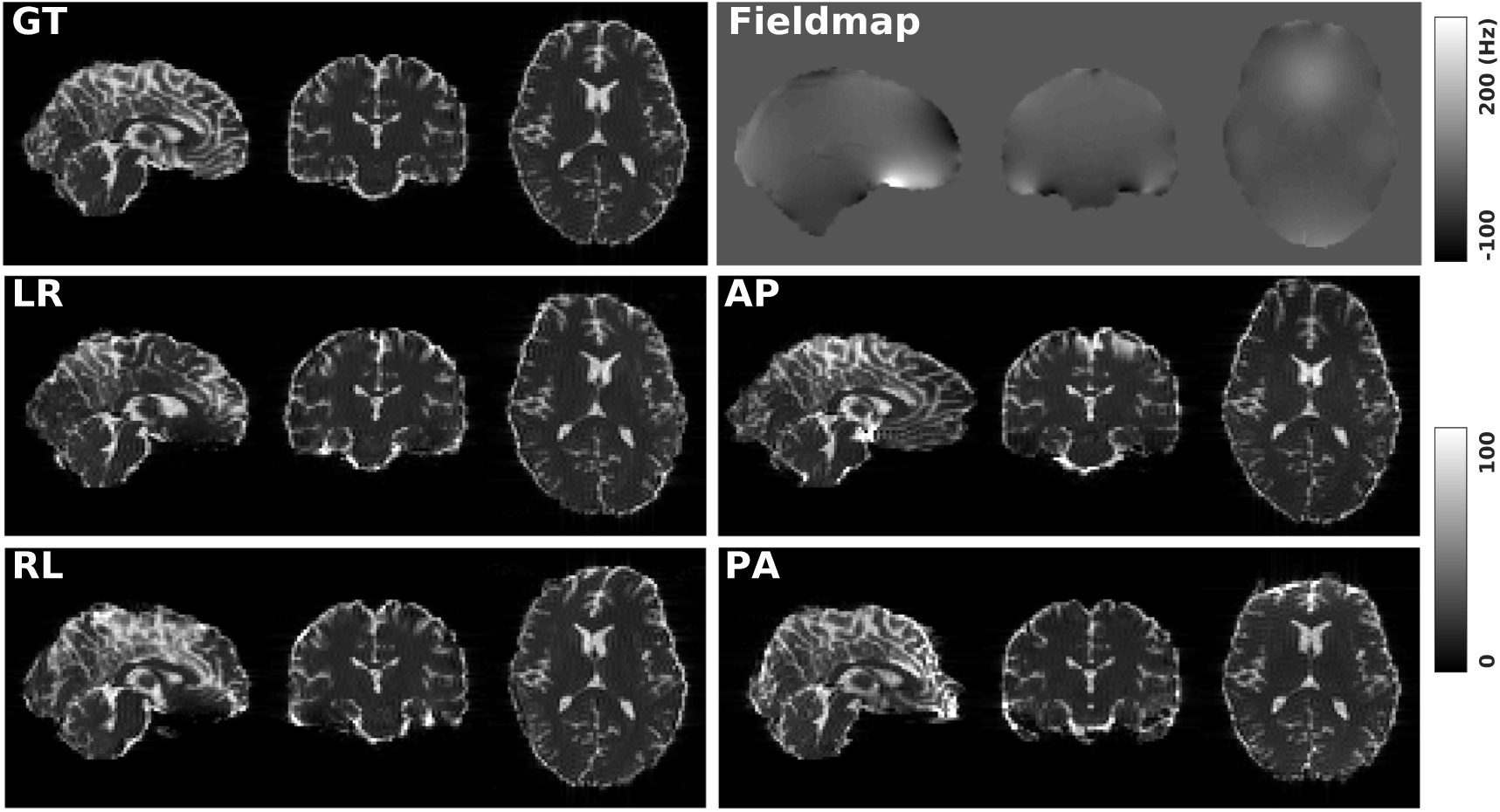
GT: simulated diffusion data (without distortions) for HCP subject 100206. Fieldmap: the real fieldmap used to simulate the distortions. LR: simulated diffusion data with LR distortion. RL: simulated diffusion data with RL distortion. AP: simulated diffusion data with AP distortion. PA: simulated diffusion data with PA distortion.

**FIGURE 2.**
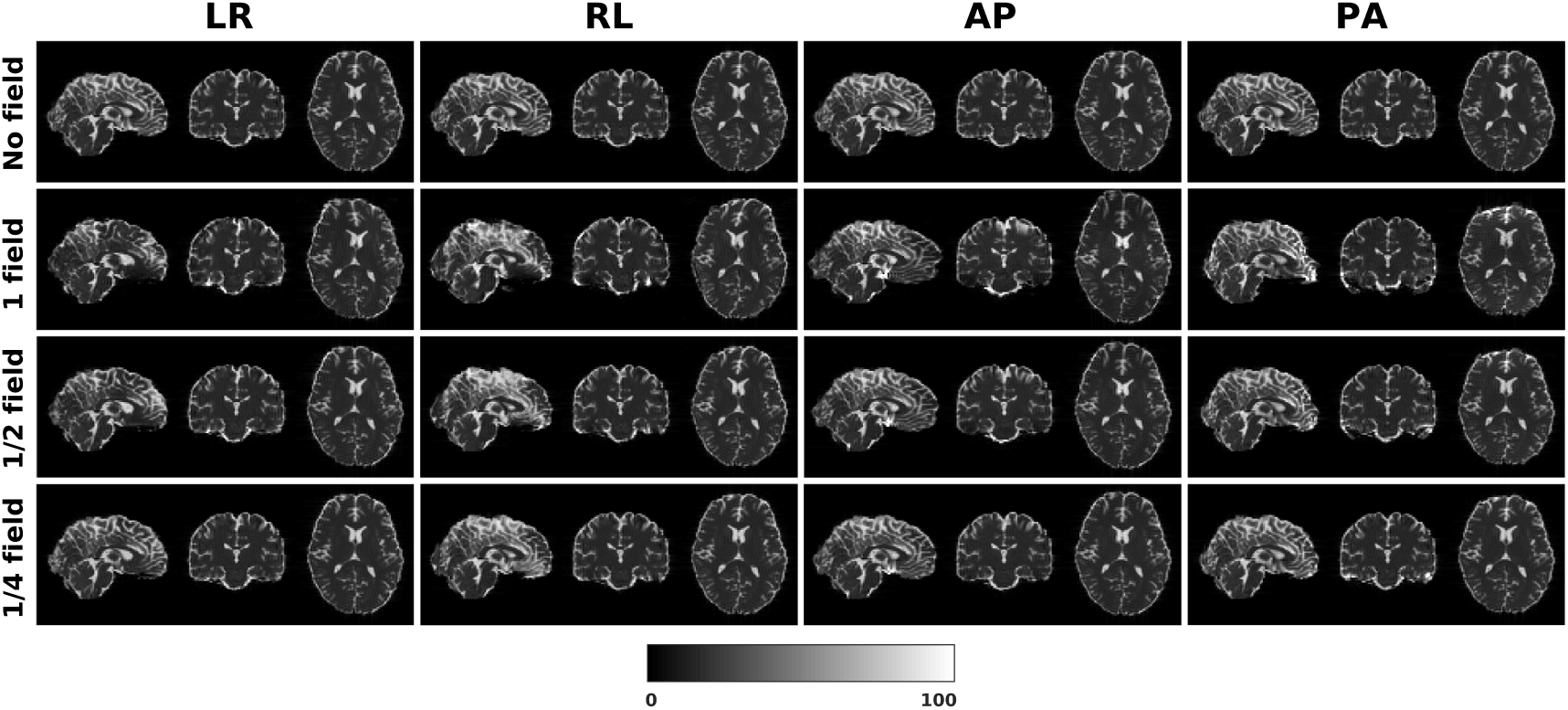
Three levels of susceptibility distortion were simulated. 1 field is the original fieldmap. 1/2 field is the original fieldmap divided by 2. 1/4 field is the original fieldmap divided by 4.

#### 4.1.1 Direct metric

In this section, we present the results for the direct metric, including the error maps of *b*_0_, FA, MD, principal eigenvalues and principal eigenvectors. Figure 3 shows the corrected *b*_0_ images using six different methods, along with error maps obtained by calculating the difference compared to the ground truth images. Correction was carried out for LRRL and APPA pairs, respectively, and we used the corrected LR and AP images as the results for the two pairs. *aDC* and *aBMDC* show large errors for edge voxels. *DR-BUDDI, EPIC, HySCO* and *TOPUP* produced very small errors for both LR and AP cases, mainly along edges.

**FIGURE 3.**
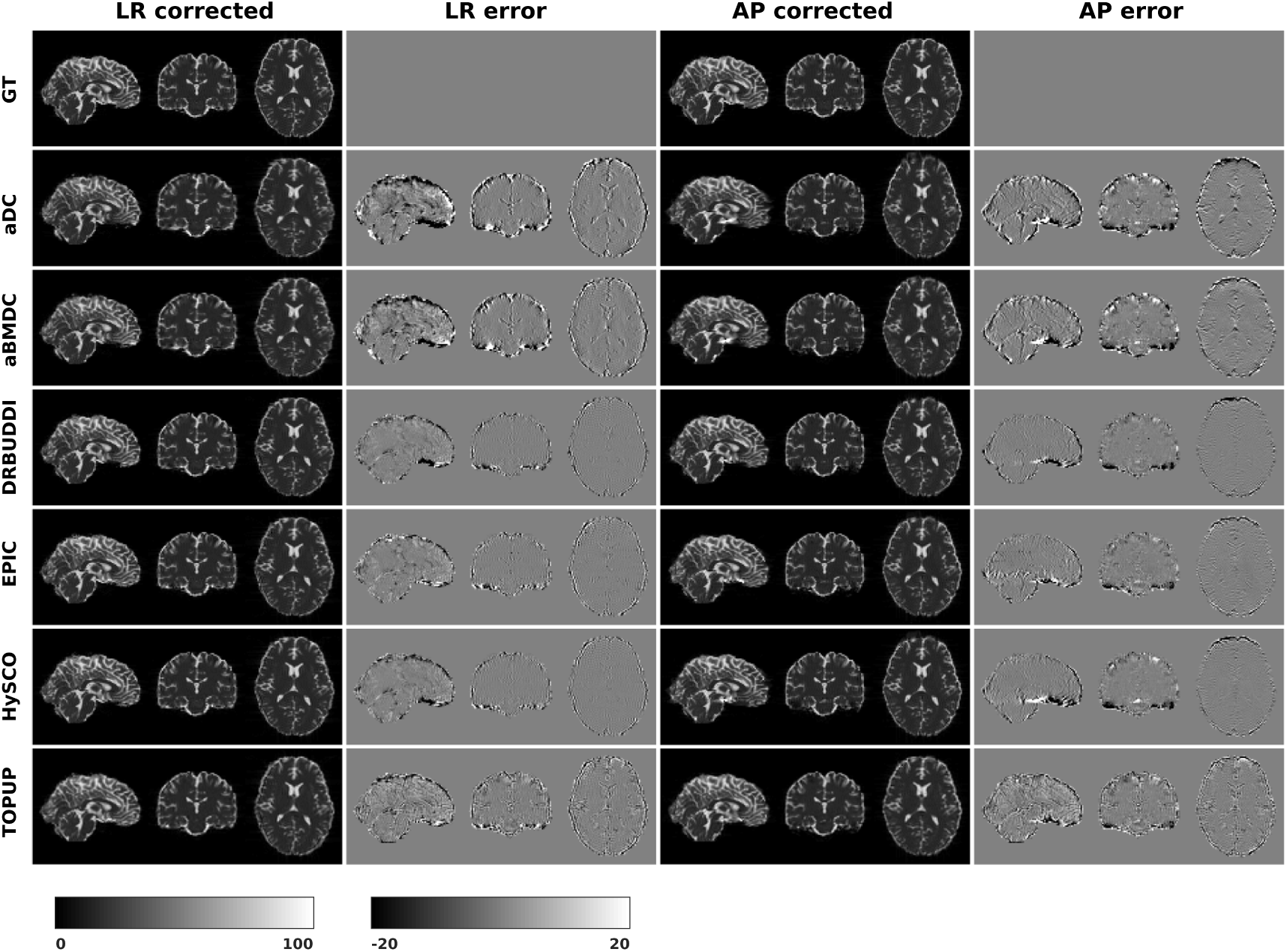
Corrected *b*_0_ volumes for simulated HCP Subject 100206 using the six methods. Error maps were obtained by calculating the difference compared to the ground truth. Correction was carried out for LRRL and APPA pairs, respectively.

Figure 4 shows the error maps using the six methods for the three levels of distortion. Correction was carried out for LRRL and APPA pairs, respectively. In addition to visual inspection, we computed the mean absolute error (MAE) and the mean squared error (MSE) within the brain and their standard deviations (error bars) for five simulated HCP subjects using the six methods, as shown in Figure 5. The results confirmed what we observed in Figure 4 and quantitatively demonstrates the accuracy and robustness of the six methods. The MAE and MSE decreased with decreasing inhomogeneity field strength, aligned with our predictions. Similarly, Figures 6 to 9 show the error maps of FA, MD, principal eigenvalues and principal eigenvectors after diffusion tensor fitting of the corrected data. The error map of principal eigenvectors was obtained by calculating the angle (in degree) between the principal eigenvector and its ground truth. All error maps are quantitatively summarized in Table 4.

**TABLE 4.** MAE and MSE of *b*_0_, FA, MD, principal eigenvalues and principal eigenvectors using the six methods. MAE and MSE are averaged across five simulated HCP subjects. For each row, the method with the best performance are in bold.

| Tool \ Metric | aDC | aBMDC | DR-BUDDI | EPIC | HySCO | TOPUP |
| --- | --- | --- | --- | --- | --- | --- |
| $b_0$ MAE, LR | 4.675 | 4.242 | **2.582** | 3.198 | 2.680 | 3.499 |
| $b_0$ MAE, AP | 4.470 | 4.029 | **2.449** | 2.698 | 2.683 | 3.369 |
| FA MAE, LR ($\times 10^{-2}$) | 2.719 | 2.698 | **1.700** | 3.143 | 3.326 | 1.923 |
| FA MAE, AP ($\times 10^{-2}$) | 2.550 | 2.577 | **1.664** | 2.454 | 3.540 | 1.829 |
| MD MAE, LR ($\times 10^{-5}$) | 3.371 | 3.224 | **2.535** | 3.588 | 3.454 | 2.584 |
| MD MAE, AP ($\times 10^{-5}$) | 3.346 | 3.197 | **2.555** | 3.070 | 3.653 | 2.587 |
| $\lambda_1$ MAE, LR ($\times 10^{-5}$) | 4.043 | 3.914 | **2.954** | 4.272 | 4.785 | 3.057 |
| $\lambda_1$ MAE, AP ($\times 10^{-5}$) | 3.926 | 3.815 | **2.959** | 3.639 | 4.985 | 3.019 |
| $v_1$ MAE, LR ($\times 10$) | 1.045 | 1.034 | **0.728** | 1.196 | 1.550 | 0.776 |
| $v_1$ MAE, AP ($\times 10$) | 1.037 | 1.032 | **0.750** | 1.011 | 1.640 | 0.781 |

**FIGURE 4.**
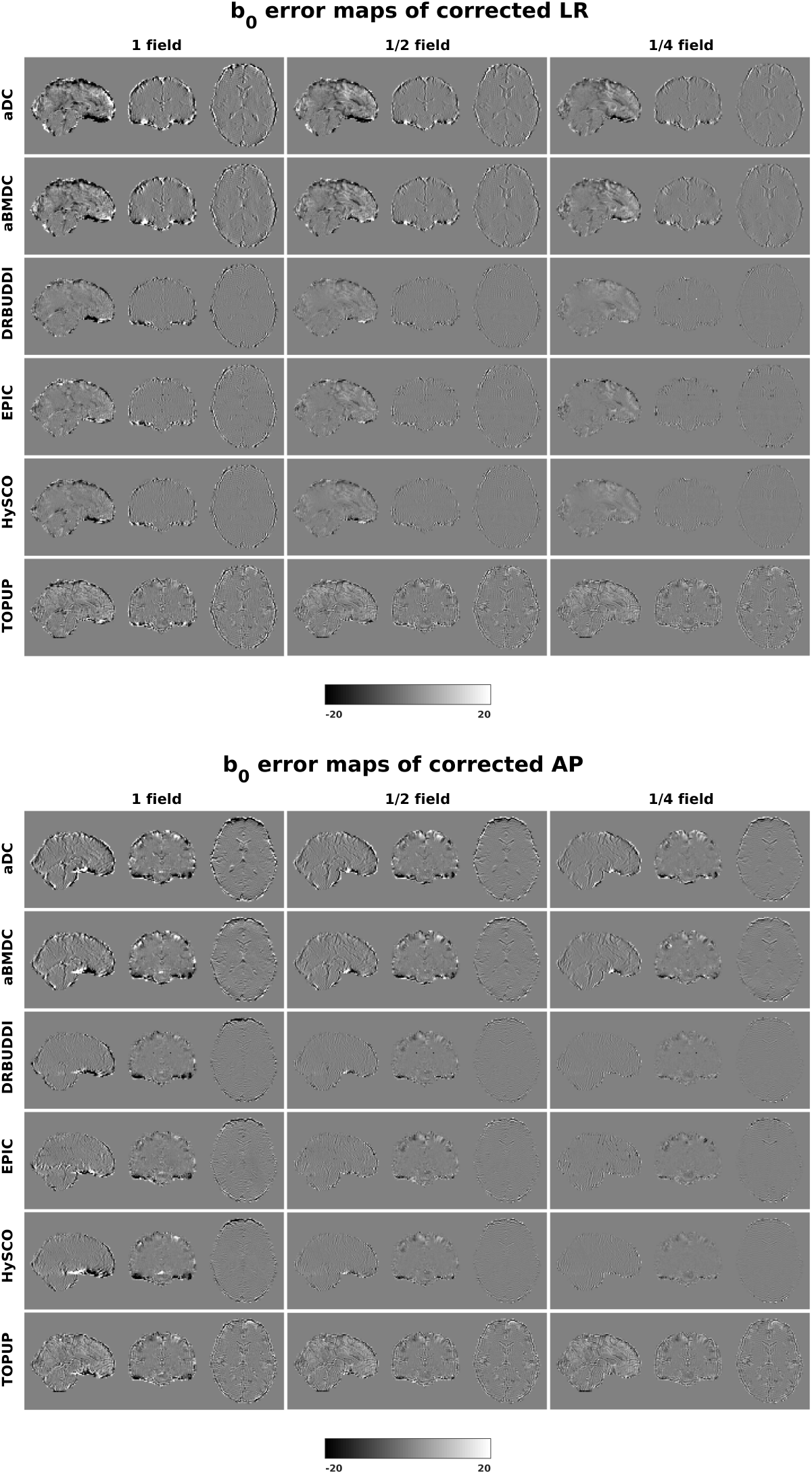
*b*_0_ error maps for simulated HCP Subject 100206 using the six methods. Three levels of susceptibility distortion were simulated. Correction was carried out for LRRL (left) and APPA (right) pairs, respectively. Three levels of susceptibility distortion were simulated.

**FIGURE 5.**
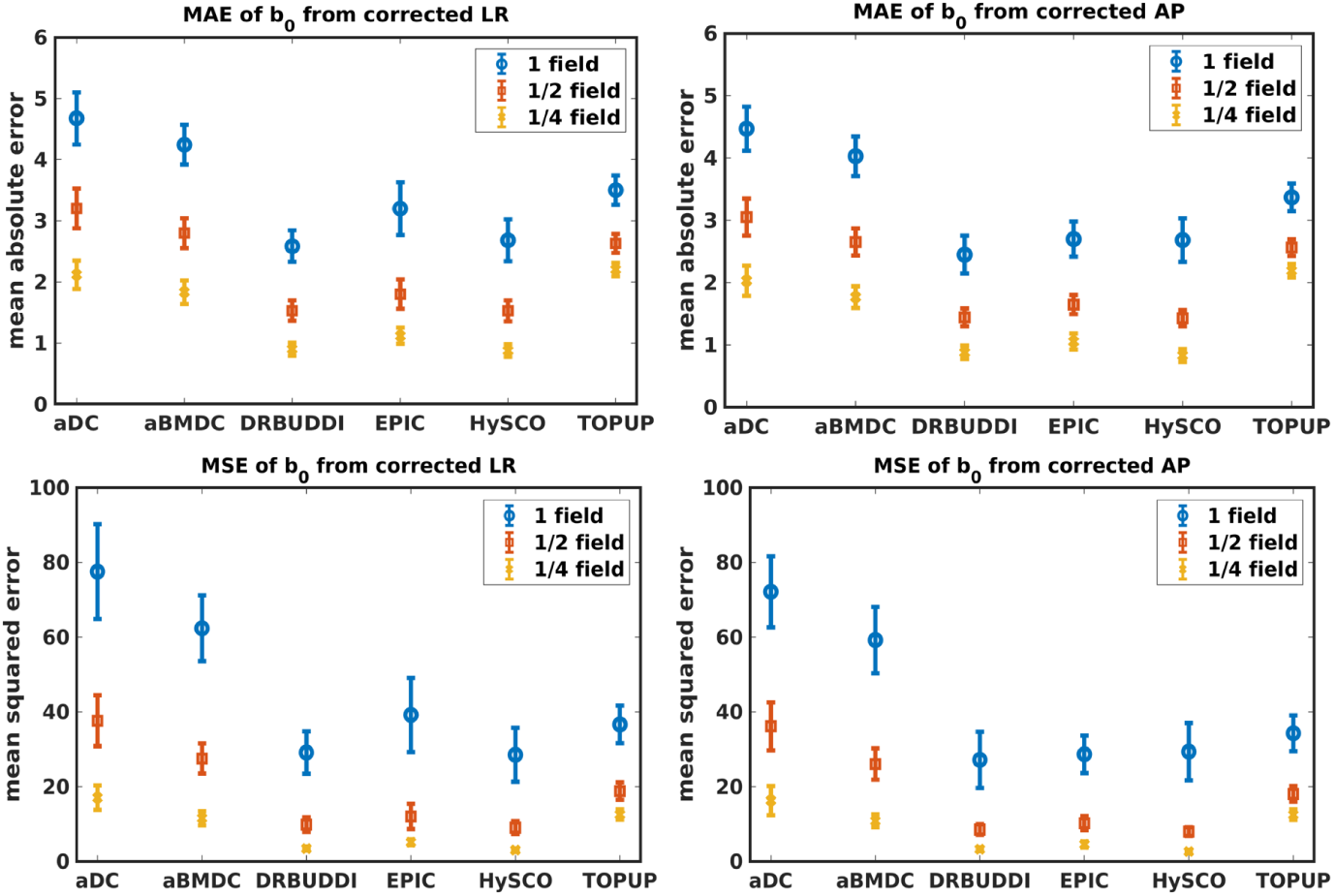
MAE (top) and MSE (bottom) of *b*_0_ for five simulated HCP subjects using the six methods. The error bars represent standard deviation over subjects. Correction was carried out for LRRL (left) and APPA (right) pairs, respectively. Three levels of susceptibility distortion were simulated.

**FIGURE 6.**
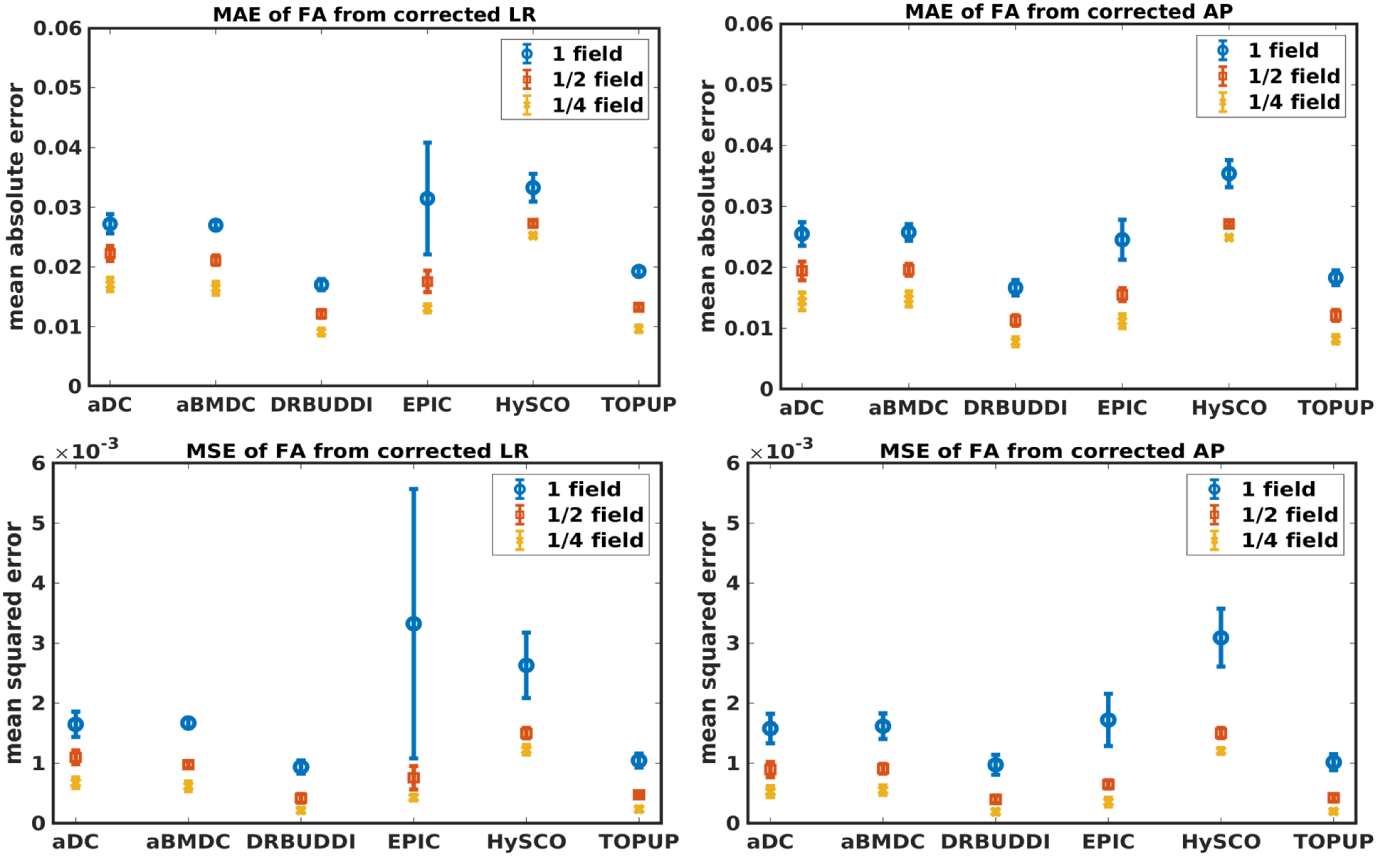
MAE (top) and MSE (bottom) of FA for five simulated HCP subjects using the six methods. The error bars represent standard deviation over subjects. Correction was carried out for LRRL (left) and APPA (right) pairs, respectively. Three levels of susceptibility distortion were simulated.

#### 4.1.2 Indirect metric

In this section, we present the results for the indirect metric, including *b*_0_ difference maps and the FA standard deviation over the corrected LR, RL, AP and PA data. To investigate the suitability of LRRL-APPA differences as an indirect metric we plot the corrected LR and AP data, and their differences, as shown in Figure 10. Ideally, the corrected LR and the corrected AP would be identical. The whole-brain mean absolute difference (MAD) and mean squared difference (MSD) were computed for every correction method, as shown in Figure 11. The results show larger MAD and MSD for *aDC* and *aBMDC* compared to the other four methods *DR-BUDDI, EPIC, HySCO* and *TOPUP*. The results are not completely consistent with the previous results in Figure 5 for the direct metric. These results demonstrate that the indirect metric (difference maps) shows a different ordering of correction performance compared to the direct metric (*b*_0_ error maps).

**FIGURE 7.**
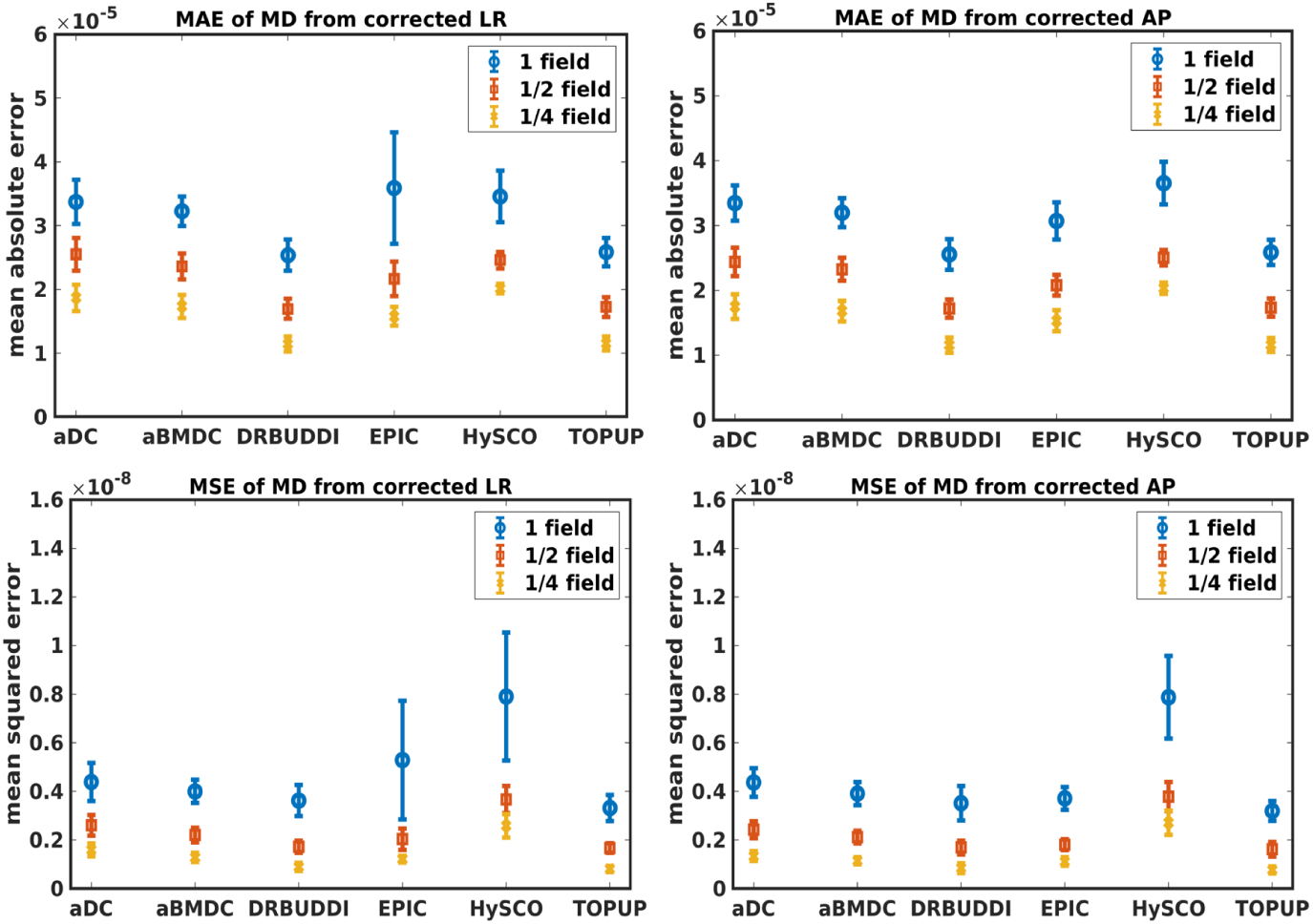
MAE (top) and MSE (bottom) of MD for five simulated HCP subjects using the six methods. The error bars represent standard deviation over subjects. Correction was carried out for LRRL (left) and APPA (right) pairs, respectively. Three levels of susceptibility distortion were simulated.

**FIGURE 8.**
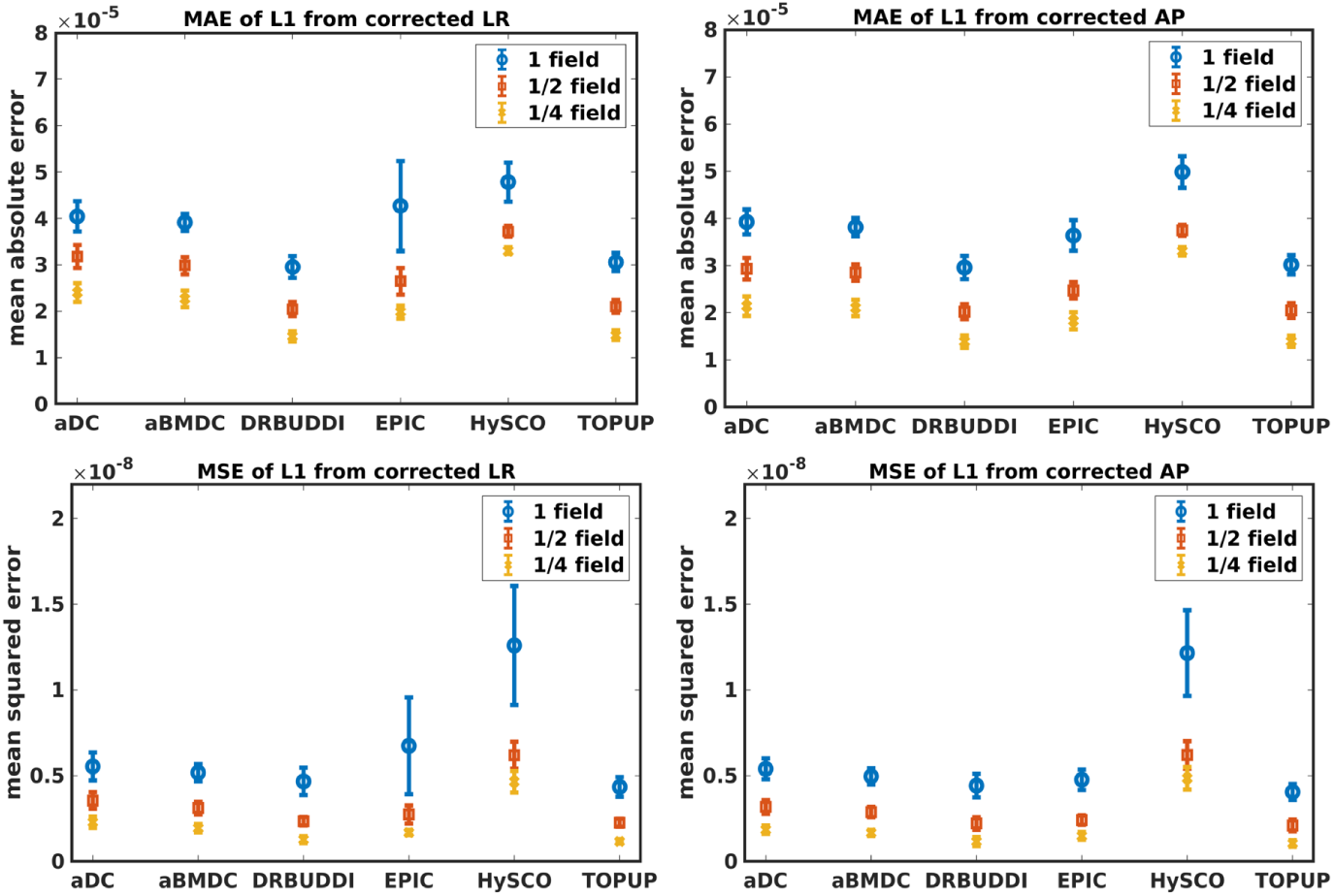
MAE (top) and MSE (bottom) of principal eigenvalues for five simulated HCP subjects using the six methods. The error bars represent standard deviation over subjects. Correction was carried out for LRRL (left) and APPA (right) pairs, respectively. Three levels of susceptibility distortion were simulated.

**FIGURE 9.**
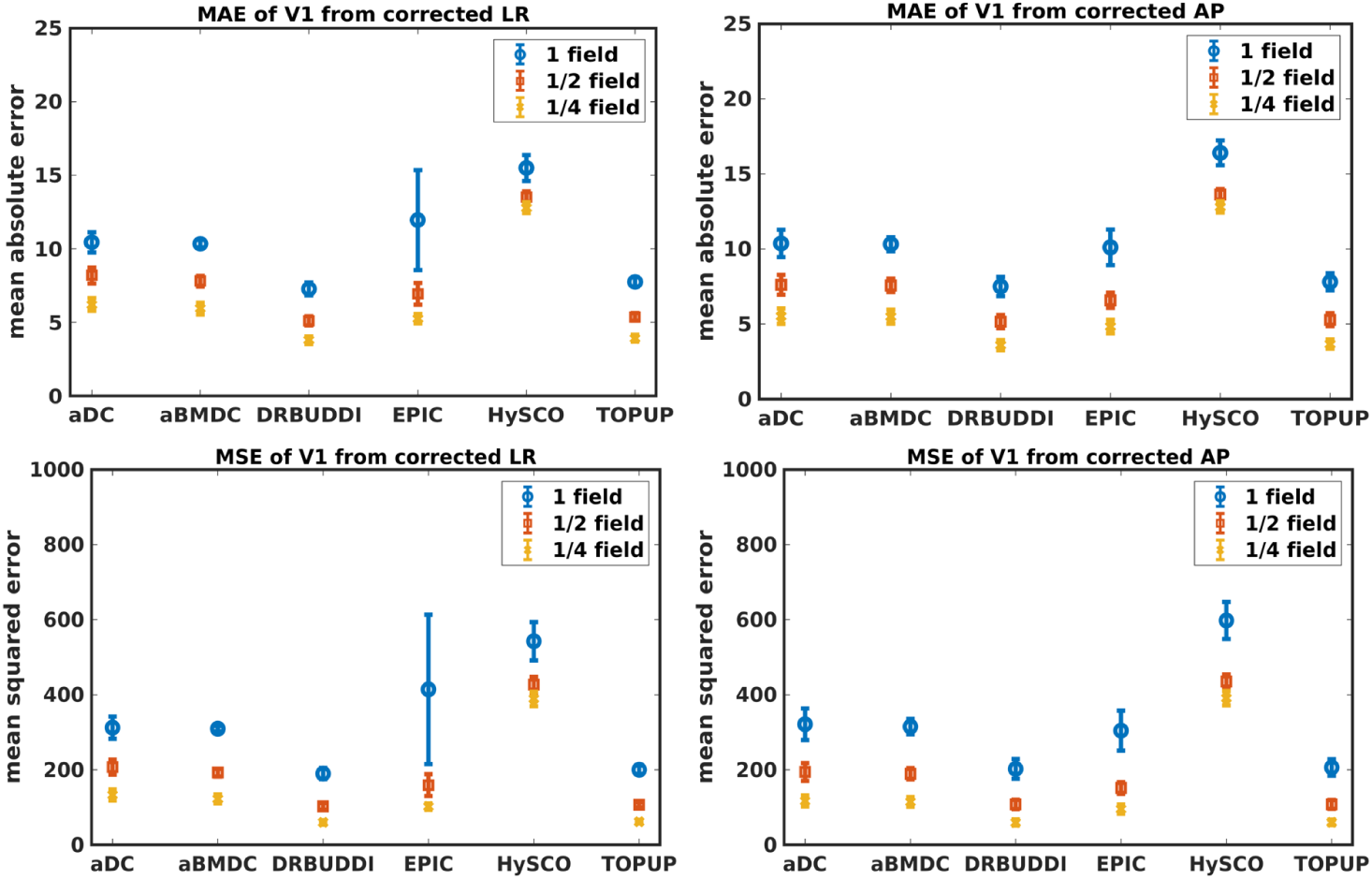
MAE (top) and MSE (bottom) of principal eigenvectors for five simulated HCP subjects using the six methods. The error bars represent standard deviation over subjects. Correction was carried out for LRRL (left) and APPA (right) pairs, respectively. Three levels of susceptibility distortion were simulated.

**FIGURE 10.**
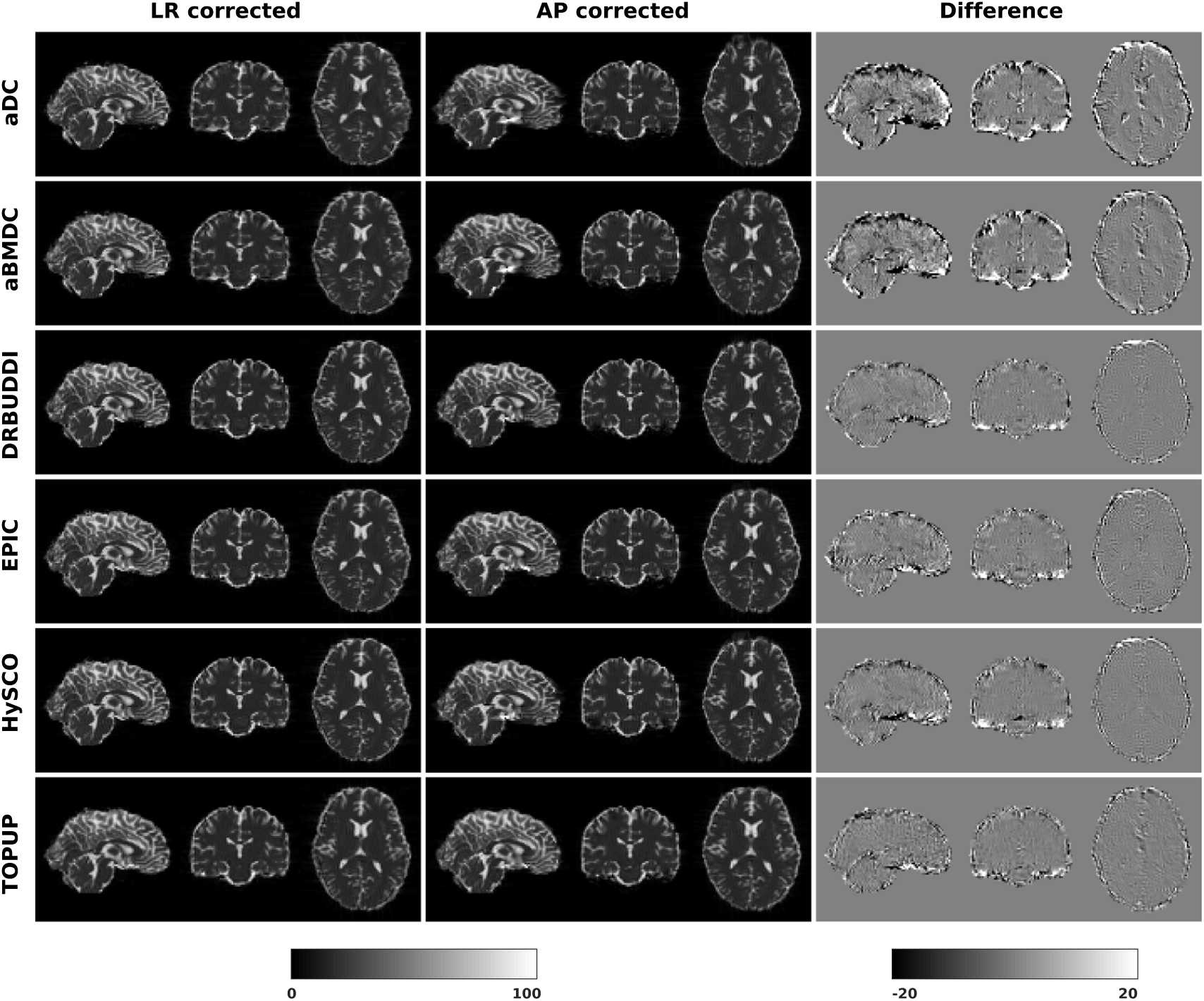
Difference between corrected *b*_0_ volumes from LR and AP for simulated HCP Subject 100206 using the six methods.

**FIGURE 11.**
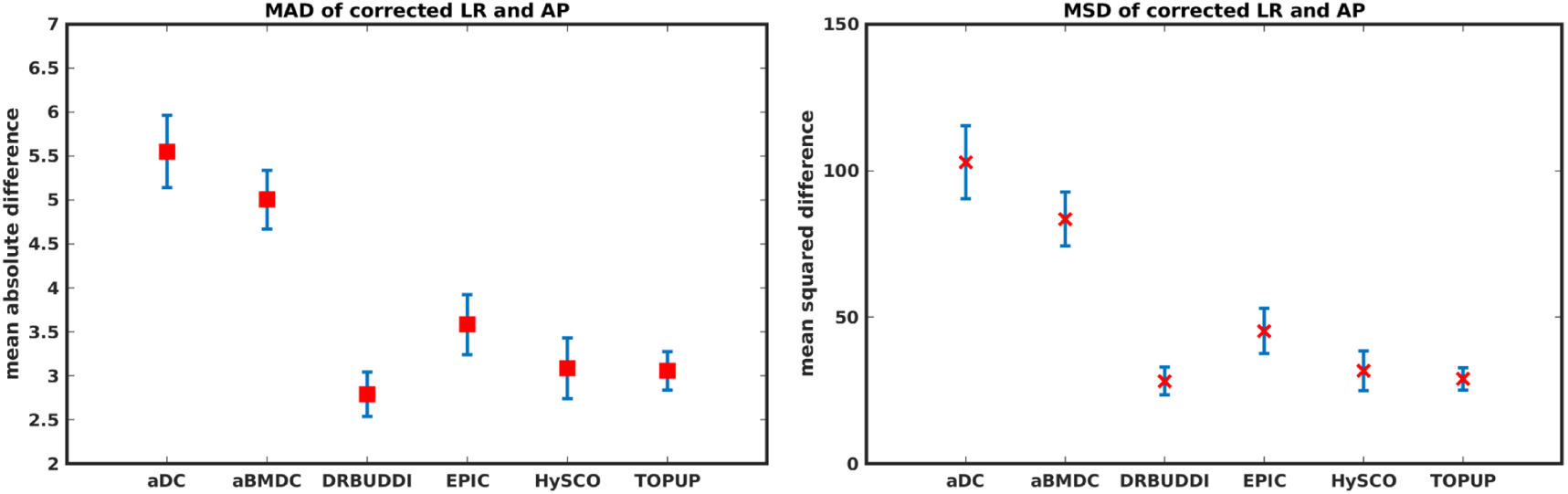
MAD and MSD between corrected LR and AP for five simulated HCP subjects using the six methods. The error bars represent the standard deviation over subjects. *DR-BUDDI* provides the smallest difference between the two corrections.

To investigate the suitability of FA standard deviation as an indirect metric we plot the FA standard deviation over the corrected LR, RL, AP and PA data, as shown in Figure 12. Ideally, the corrected LR, RL, AP and PA would be identical, which would result in zero FA standard deviation. The whole-brain mean of the standard deviation was computed for every correction method, as shown in Figure 13. The results show the largest mean FA standard deviation for *EPIC. DR-BUDDI* and *TOPUP* produced very small FA standard deviation over LR, RL, AP and PA data, and the standard deviation of this metric is also low across subjects. The indirect metric (FA standard deviation) served as a good indication for correction performance compared to the direct metric (FA error maps) as shown in Figure 6.

**FIGURE 12.**
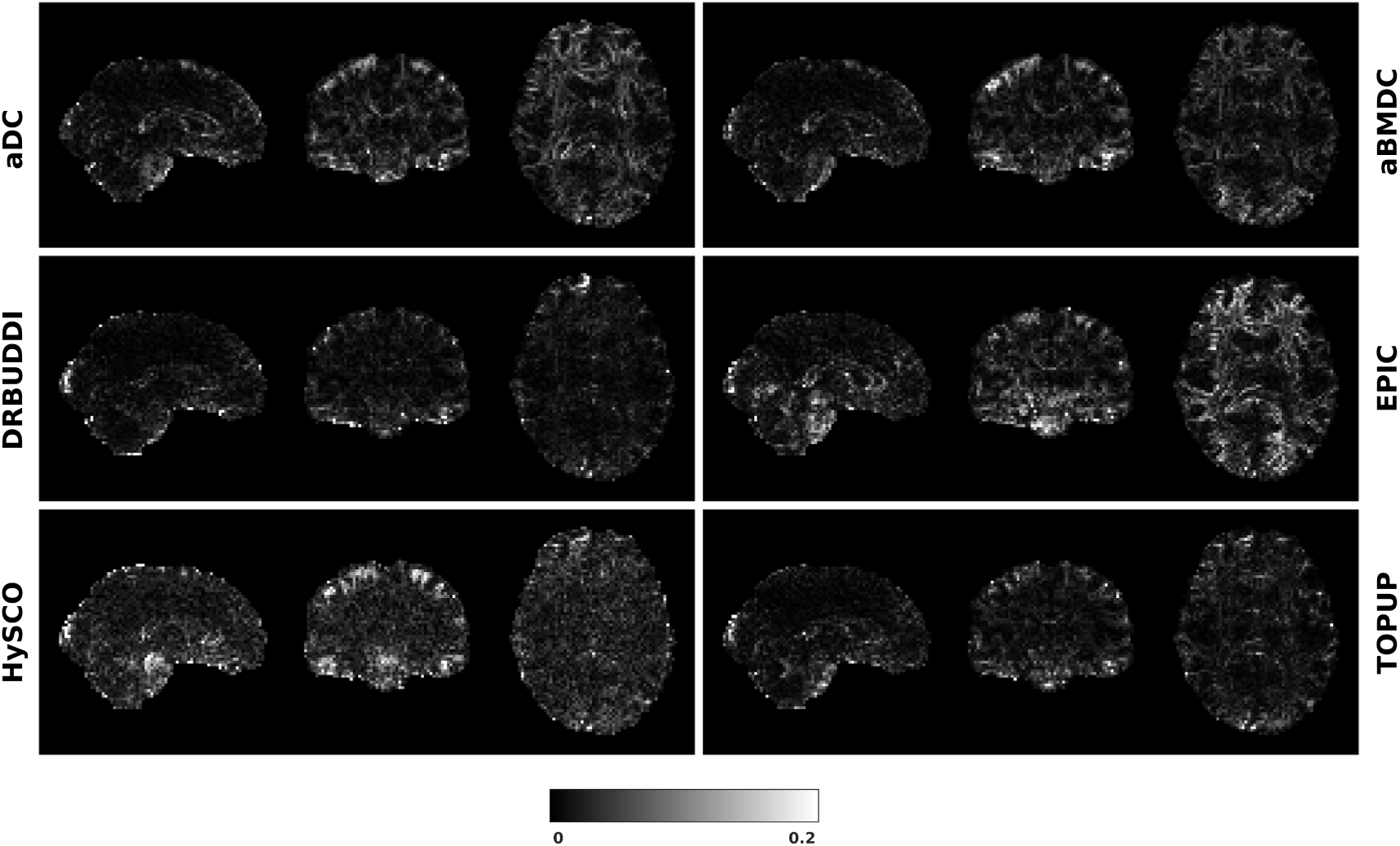
FA standard deviation over the corrected LR, RL, AP and PA data for simulated HCP Subject 100206 using the six methods.

**FIGURE 13.**
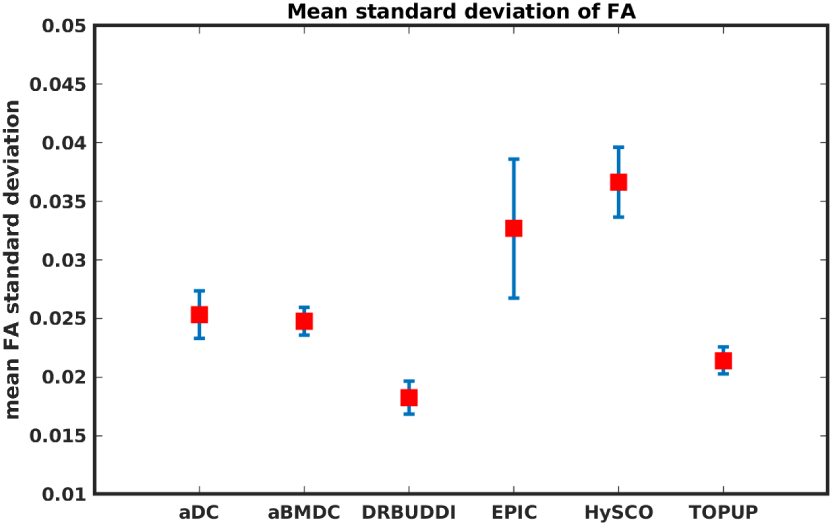
Mean FA standard deviation over LR, RL, AP and PA data for five simulated HCP subjects using the six methods. The error bars represent standard deviation of this metric across subjects. *DR-BUDDI* produces the smallest standard deviation.

### 4.2 Real data

Figure 14 shows the distorted diffusion data from a dHCP subject scanned with four different PE directions. The distortions are more pronounced in regions close to tissue-air interfaces, such as the frontal poles and the temporal lobes near the petrous bone. To evaluate the six methods, we computed the difference between the corrected LR and AP for the dHCP data, as shown in Figure 15. Higher difference values are visible in regions most affected by magnetic susceptibility variations such as the boundary regions of the brain. Figure 16 reports the whole-brain mean of the difference, for the 40 processed subjects. *DR-BUDDI, EPIC, HySCO* and *TOPUP* performed almost equally well, which resembled the results for simulated data given in Figure 11.

**FIGURE 14.**
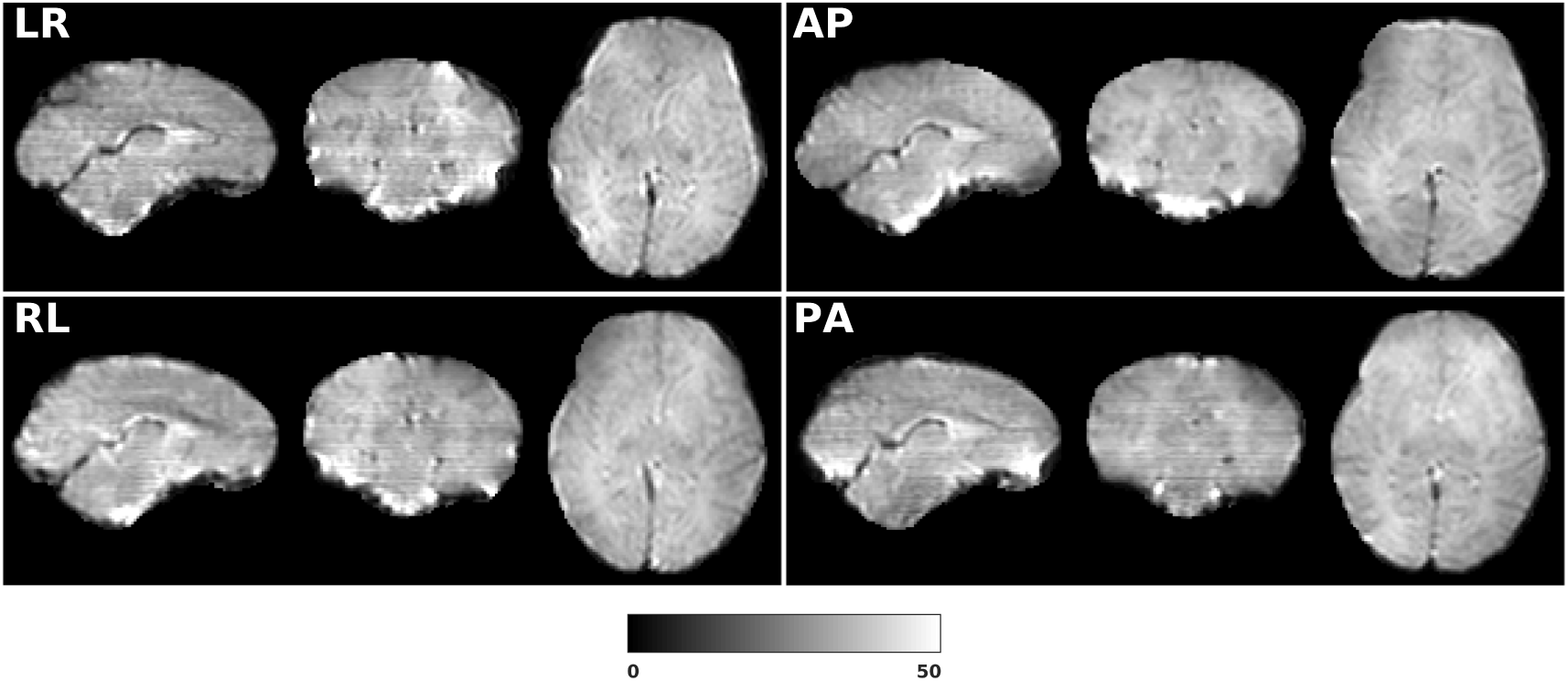
LRRL and APPA pairs for dHCP Subject CC00069XX12.

**FIGURE 15.**
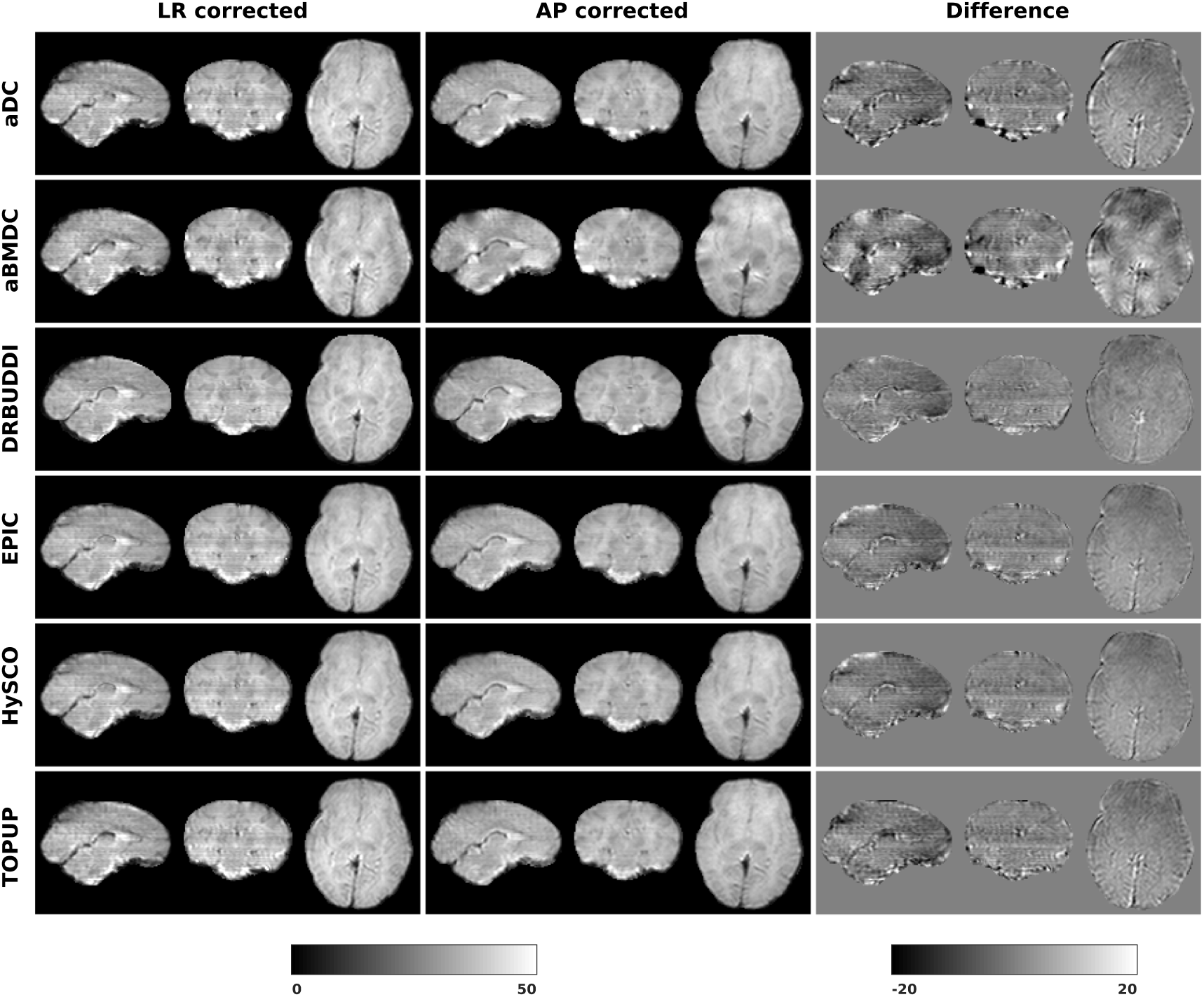
Difference between corrected data from LRRL and APPA pairs for dHCP Subject CC00069XX12 using the six methods.

**FIGURE 16.**
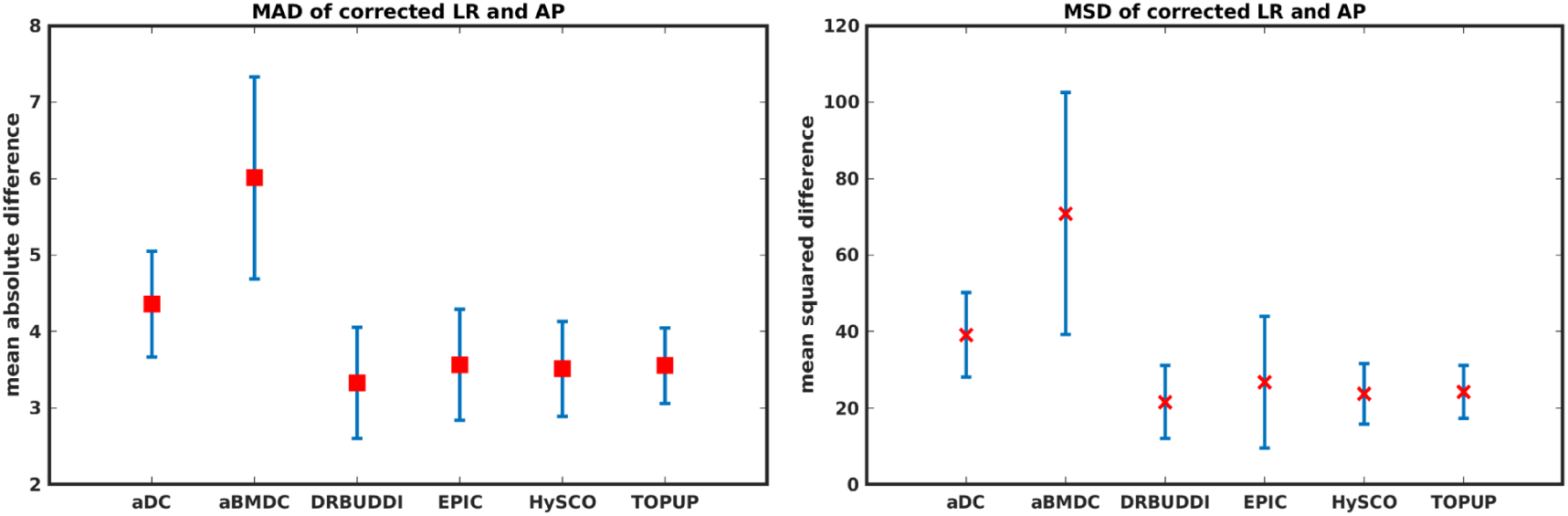
MAD and MSD between corrected data from LRRL and APPA pairs for 40 dHCP subjects using the six methods. The error bars represent the standard deviation over subjects. In all cases *DR-BUDDI* produces the smallest difference.

### 4.3 Processing time

The processing time of one simulated HCP dataset for the six softwares are 2.8 (*aDC*), 35.5 (*aBMDC*), 153.3 (*DR-BUDDI* without its pre-step *DIFFPREP*), 38.0 (*EPIC*), 8.1 (*HySCO*) and 195.7 (*TOPUP*) seconds, respectively. Please note that *aBMDC, DR-BUDDI, EPIC* and *HySCO* use several CPU threads to speedup the processing. We used a computer with 32 GB RAM and an Intel(R) Xeon(R) Silver 4114 2.20 GHz CPU (containing 10 cores, which can run 20 threads in parallel).

## 5 DISCUSSION

## 5.1 Discussion

In this paper, we used both simulated and real data to evaluate six phase encoding based methods for correcting susceptibility distortions. This work is important given that phase encoding based methods have been demonstrated to outperform the other two classes of approaches, and are very frequently used in diffusion data analysis pipelines. It is thus essential to carefully evaluate phase encoding based correction techniques and their limitations. By this work we aim to answer the following two questions. Which method of the six provides the best distortion correction? Are the indirect metrics suitable for measuring the distortion correction performance?

The error map directly measures the ability to correctly recover distortion-free data for different methods. Based on our experiments, we found that *DR-BUDDI, EPIC, HySCO* and *TOPUP* were generally superior to the other two methods in achieving better performance for the error map of *b*_0_, see Figure 5. We then applied the transformation field obtained from *b*_0_ images to diffusion weighted volumes and calculated FA, MD, eigenvalues and eigenvectors after diffusion tensor fitting. The error maps of these four metrics demonstrated that *DR-BUDDI* and *TOPUP* provided the most accurate and robust distortion corrections among the six methods. *EPIC* and *HySCO* performed well in correcting *b*_0_ images but produced poor corrections for diffusion weighted volumes, and these two methods produced rather large errors in terms of the four diffusion tensor metrics as shown in Figures 6 to 9. Therefore, the error map of *b*_0_ should be interpreted together with the error maps of diffusion metrics for a better evaluation of the correction quality. It is notable that *EPIC* showed large errors and very large variance of performance over subjects for the LRRL case when the inhomogeneity field strength was 1. The reason might be that *EPIC* was originally designed for the APPA case and there is no method referred to in any of the *EPIC* documentation for choosing an alternate phase direction than the *y*-direction (or the LRRL direction with regards to this study). With the phase encode direction chosen in the APPA dimension, susceptibility distortion is manifested as stretching or compression of the image in the APPA direction, which can be more desirable than asymmetric distortion in the LRRL direction (Glover et al., 2012). This might explain that EPIC was designed only for the APPA case. It is indeed often the case that the phase encoding direction is chosen to be the *y*-direction, however there are many images where this is not the case (Hughes et al., 2017; Kemper et al., 2015). *TOPUP* was found to work only for data volumes with even size on *x, y* and *z* directions since the default configuration file (*b02b0.cnf*) performs a factor of 2 subsampling. There are two solutions suggested to eliminate this problem; either cropping or adding dummy data to the 3D volume.

Indirect metrics are often used to evaluate distortion correction for real data due to the absence of ground truth. We investigated the ability of two indirect metrics to measure the correction quality. We validated two of the most promising indirect metrics for correction quality, i.e., the difference between corrected LR and AP data, and the FA standard deviation over the corrected LR, RL, AP and PA data. The first indirect metric, as shown in Figures 11 and 16, roughly confirmed what we observed for the error map of *b*_0_ in Figure 5, although the ordering of correction quality is slightly different. Similarly, the FA standard deviation over the corrected LR, RL, AP and PA data, as shown in Figures 13, confirmed what we observed for the error map of FA in Figures 6. We therefore suggest that indirect metrics must be interpreted cautiously, and the two indirect metrics should be interpreted together as a measure of distortion correction quality.

## 5.2 Limitations

Before comparing the quality of the corrections provided by each of these tools, it is important to note that each tool has its customizable parameters. The default parameter settings were used in this work, as it was reasoned that this would be representative of the way the tools were most often used. It was also reasoned that changing default parameters or influencing the inputs prior to correction would lead to a less fair comparison. It should be noted that we used different *DR-BUDDI* settings for simulated data and real data, because the default pipeline is designed for real data with different types of distortions, but our simulated data only contain susceptibility distortions. It is therefore possible that the corrections in this work may not represent the best corrections attainable by use of these tools.

## Acknowledgements

This research was supported by the Swedish Research Council (grant 2017-04889), Linköping University Center for Industrial Information Technology (CENIIT) and the ITEA3 / VINNOVA funded project “Intelligence based iMprovement of Personalized treatment And Clinical workflow supporT” (IMPACT). We used the data provided by the Human Connectome Project, WU-Minn Consortium (Principal Investigators: David Van Essen and Kamil Ugurbil; 1U54MH091657) funded by the 16 NIH Institutes and Centers that support the NIH Blueprint for Neuroscience Research; and by the McDonnell Center for Systems Neuroscience at Washington University. Part of these results were obtained using data made available from the Developing Human Connectome Project funded by the European Research Council under the European Union’s Seventh Framework Programme (FP/2007-2013) / ERC Grant Agreement no. [319456]. The authors would like to thank Mark Graham for providing help with the POSSUM diffusion MRI simulator. The authors would also like to thank Renaud Hedouin, Olivier Commowick, Mustafa Okan Irfanoglu, Ivar Thokle Hovden, Lars Ruthotto and Jesper Andersson for providing comments on the usage of their softwares.

## CONFLICT OF INTEREST

The authors declare no conflict of interest.

## Footnotes

1 Data collection and sharing for this project was provided by the Human Connectome Project (U01-MH93765) (HCP; Principal Investigators: Bruce Rosen, M.D., Ph.D., Arthur W. Toga, Ph.D., Van J.Weeden, MD). HCP funding was provided by the National Institute of Dental and Craniofacial Research (NIDCR), the National Institute of Mental Health (NIMH), and the National Institute of Neurological Disorders and Stroke (NINDS). HCP data are disseminated by the Laboratory of Neuro Imaging at the University of Southern California.

2 https://github.com/xuagu37/SusceptibilityDistortionCorrection

## Notes

#### Summary of Updates

Updated results for DR-BUDDI, added summary table

